# Stable isotope insights into artificial reef effects of floating offshore energy structures in Norwegian North Sea codfishes

**DOI:** 10.1101/2025.10.20.683384

**Authors:** Adam Jon Andrews, Steven Brooks

## Abstract

Offshore energy structures introduce hard substrate to soft substrate-dominant habitats and may act as artificial reefs providing shelter and food to aggregating fish. In the Northeast Atlantic, knowledge on these effects is limited to shallow habitats in the southern North Sea. Given that effects may be misinterpreted as ‘nature positive’ contributions, or underestimated and impacting ecosystem services like fisheries, this data-gap hinders management. This is especially problematic for the rapid developments of floating offshore wind farms (OWFs), and decommissioning of oil and gas (O&G) platforms in deep (>100 m) habitats of Norway, the U.K. and Ireland. In this study, we analysed the stable isotopic composition of muscle and liver and the condition of three codfishes of commercial importance (saithe; *Pollachius virens*, tusk; *Brosme brosme* and ling; *Molva molva*) at a floating OWF and two floating O&G platforms off Norway to evaluate how codfish diet and habitat use may be altered by the structures. We find that differences in carbon, nitrogen and sulphur stable isotopes between offshore energy sites and control sites were lower for liver measurements (weeks prior to capture) than muscle (months prior to capture), indicating that codfish diet and habitat use was less impacted by offshore energy structures than longer-term natural feeding variation. Size explained some isotopic differences between sites in saithe, and condition differences between sites in tusk; suggesting that the diet and habitat use of the three species is not significantly impacted by offshore structures. However, we found evidence of lower condition in ling at Hywind Tampen OWF, corresponding to lower nitrogen isotope liver values in ling; that may indicate trade-offs in shelter and diet provision. Overall we highlight the need for further research on trophic effects of deep offshore energy structures to evaluate implications for management and conservation.

## 1. Introduction

Offshore wind farm (OWF) developments are occurring rapidly, resulting in an urgent need to assess their environmental consequences. Among them, the roles of offshore energy structures (platforms, turbines, foundations and scour protection etc.) as artificial habitats for aggregating mobile organisms like fishes, emerge as significant uncertainties–especially for understudied areas of development like deeper (>100 m) habitats and newer technologies like floating foundations (Pardo et al., 2023). So-called artificial reef effects are a phenomenon from decades of research in shallow habitats but questions remain as to what artificial reef effects exist in deeper waters, using floating technologies, and how they impact populations. Research e.g. from shallow (<50 m) southern North Sea habitats clearly shows elevated abundance and/or diversity of fishes near offshore energy structures (Fujii, 2016; Reubens et al., 2013b; Stenberg et al., 2015), even at larger scales i.e. extending kilometers away (Lawrence et al., 2024). However, it remains unclear whether food provisioning is driving these differences as suggested in plaice (*Pleuronectes platessa*, (Buyse et al., 2023)), saithe (*Pollachius virens* (Fujii, 2016)), pouting (*Trisopterus luscus*) and sculpin (*Myoxocephalus scorpioides*, (Mavraki et al., 2021; Reubens et al., 2013b)) but not commonly or consistently evident for many fishes (Fujii, 2016). This is true especially for pelagic fishes but also bentho-pelagic ones like Atlantic cod (*Gadus morhua*) (Mavraki et al., 2021; Reubens et al., 2014, 2013b).

Offshore energy structures i.e. artificial hard-bottom habitats clearly can host assemblages of fouling and sheltering benthic organisms that vary from natural soft-bottom assemblages (Coolen et al., 2020; Guerin 2009), including prey items like two-spotted gobies (*Gobiusculus flavescens*) and edible crab (*Cancer pagurus*) that can reproduce at such sites (Krone et al., 2017; van Hal et al., 2017). Fish become attracted, at least temporarily i.e. diurnally, over weeks or months (Fujii and Jamieson, 2016; Lawrence et al., 2024; Reubens et al., 2013a). However, if artificial habitats only serve to attract and aggregate fish, by definition they are not increasing the productivity of fish populations. Instead, they make fish more easily caught at site boundaries. This has consequences for management given that artificial reef effects may not be potential ‘nature positive effects’ (Pardo et al., 2023) but ‘ecological traps’, resulting in issues for fisheries sustainability and potentially further declines in overexploited fisheries (c.f. “attraction–production debate” (Grossman et al., 1997; Reubens et al., 2013b)).

Effects of offshore energy structures on fish diet and habitat use in deeper northern North Sea habitats are especially unclear due to a lack of studies. Therefore, we hypothesise that effects are limited due to: 1) overall low fish densities (Lawrence et al., 2024); 2) benthic invertebrate diversity typically decreasing with depth (van der Stap et al., 2016); and 3) that floating-foundation design in deep water provides relatively less shelter than fixed foundations in shallow waters. Stable isotope analysis (SIA) of carbon ( δ^13^C), nitrogen (δ^15^N) and sulphur (δ^34^S) offers the potential to reveal effects of structures on diet and habitat use. Similar approaches are rare (Buyse et al., 2023; Mavraki et al., 2021; Plumlee et al., 2021) but are invaluable proxies for diet and habitat use because they provide longer-time perspectives than stomach content analysis e.g. (Lawrence et al., 2024; Reubens et al., 2014); and allow study when stomach expulsion rates are high–which is often the case when sampling deep-dwelling fish (e.g. Tenningen et al. 2024). Depending on species and tissue type, isotope signatures represent diet contributions over longer time frames ca. months before capture (for muscle) or ca. several weeks (for liver) (Boecklen et al., 2011).

A range of ecological and environmental variables will affect the δ^13^C, δ^15^N and δ^34^S values of fish tissues. δ^15^N values increase with each trophic level and are thus used to estimate trophic position (Sigman et al., 2009). In contrast, δ^13^C and δ^34^S signatures pass between primary producers and consumers with low levels of fractionation. This lends them to be reliable indicators of provenance because distinct δ^13^C and δ^34^S values are generally maintained across trophic levels (Guiry, 2019; Thode, 1991). Pelagic habitats and consumers often contain lower δ^13^C than benthic and neritic ones (Amiraux et al., 2023; DeNiro and Epstein, 1978) yet pelagic δ^15^N values are often higher due to high levels of fractionation and that pelagic food webs are more complex than benthic ones. δ^34^S is sometimes useful to disentangle provenance further. For example, low δ^34^S values often reflect increased foraging on benthic or neritic prey while higher values indicate a greater degree of energy incorporated from pelagic production (Raoult et al., 2024). Distance from shore also influences δ^34^S values, i.e. food webs associated with coastal habitats such as seagrass beds, salt marshes and mudflats that incorporate low δ^34^S from anoxic marine sediments during production (Guiry et al., 2022; Szpak and Buckley, 2020; Thode, 1991).

In this study, we examine differences in stable isotope signatures of three bentho-pelagic codfishes (saithe *Pollachius virens*, tusk *Brosme brosme* and ling *Molva molva*) between well-established floating offshore oil and gas (O&G) structures (Snore), a new (2022) floating OWF i.e. Hywind Tampen, and inshore and offshore control sites in the Norwegian North Sea (Figure 1). First, we analysed stable carbon (δ^13^C), nitrogen (δ^15^N) and sulphur (δ^34^S) values to test for differences in fish diet between sites. Second, we tested for temporal differences between muscle and liver tissue that can explain site dependence. Third, we assessed explanatory variables like body size and condition.

**Figure 1.**
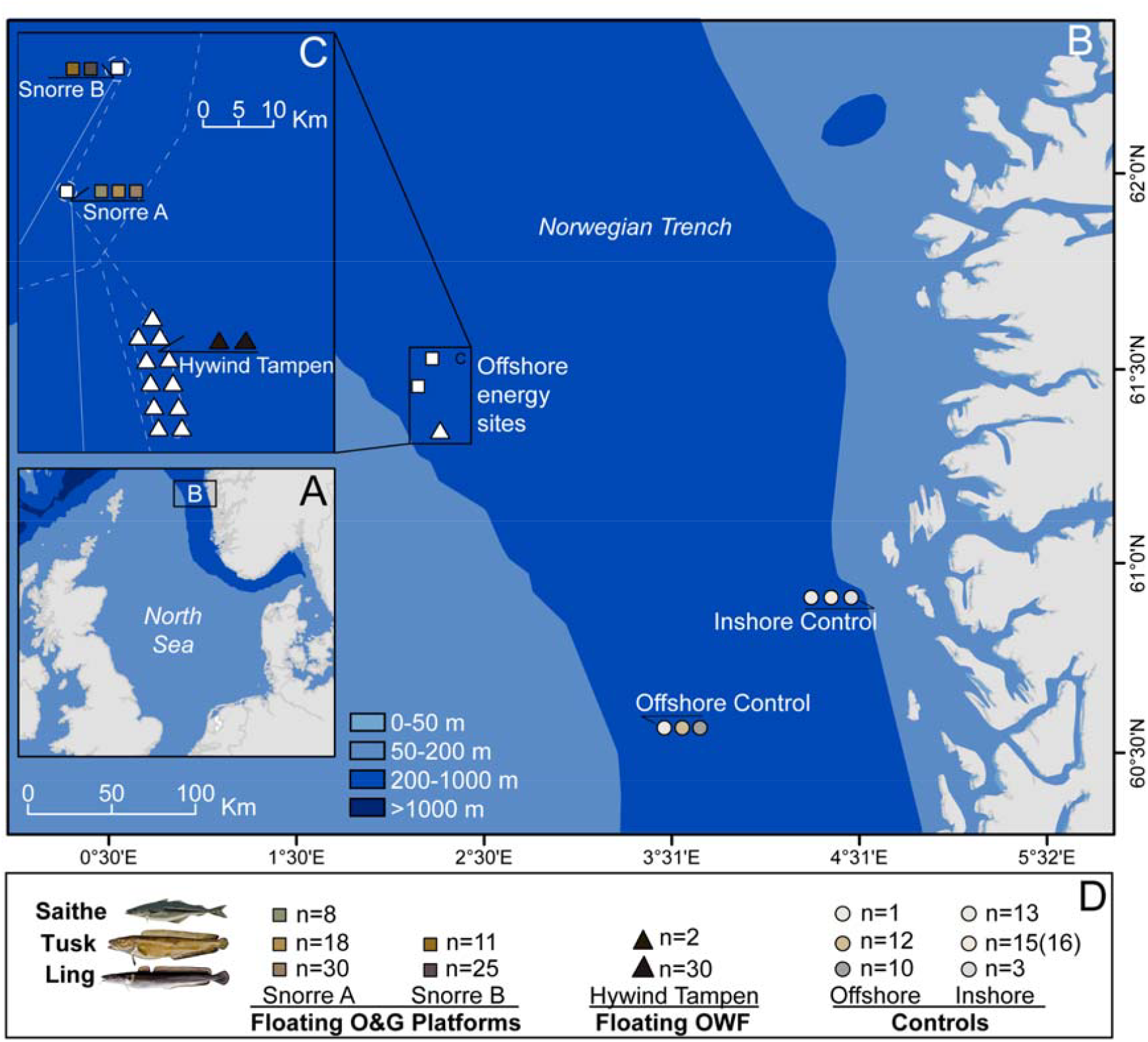
Map showing the location of the Norwegian study site in the North Sea (A), locations and distance between sample sites (B) including a detailed view of offshore energy sites (C.). Details of sampling design (D) show numbers of individuals analysed per species (illustrated cartoons); Saithe (*Pollachius virens*; green), Tusk (*Brosme brosme*; tan), and Ling (*Molva molva*; greyscale and brown). Numbers (n) are analysed measurements on muscle and liver except the number of liver samples for Inshore Control Tusk – as indicated by parentheses. Black line arrows point to sampling locations. Square shapes depict oil and gas (O&G) floating platforms, triangles depict floating offshore wind turbines and circles represent natural control sites; inshore and offshore. Cables are indicated by dashed white lines, pipelines are indicated by full white lines. Map created using ESRI ArcMap (v.10.6, https://arcgis.com).

## 2. Materials and Methods

### 2.1 Sampling and sample processing

A total of 179 individuals of three bentho-pelagic marine fish—saithe (Pollachius virens, n=22), tusk (Brosme brosme, n=59), and ling (Molva molva, n=98)—were caught off Norway between during two sampling campaigns, 17th - 27th March and 27th April to 3rd May 2024 as part of the Water Column Monitoring 2024 programme coordinated by the energy developer Equinor ASA (Figure 1). Fish were caught by hook-and-line using mackerel-baited lures at five sites: two O&G platforms, Snorre A and Snorre B (at approximately 310 m depth, within 500 m of the platforms); one OWF, Hywind Tampen (at approximately 290 m depth, within 500 m of the turbines); and two control sites: one inshore control (at approximately 100-140 m depth) outside Fensfjorden, and one offshore control (at approximately 300 m depth) in the Oseberg region (Figure 1). The offshore control site is approximately 30 km away from O&G activities, Hywind Tampen is ca. 10 km away from Snorre A, and Snorre A and B are ca. 7 km apart. Total length (cm), total weight (g), liver weight (g), and stomach weight (g) were recorded for each individual to the nearest cm or gram, respectively. Fulton’s Condition Index (K = 100 × weight / length^3^) was calculated from these measurements. White muscle (excluding skin) and liver samples were collected from individuals, snap frozen in liquid nitrogen and stored frozen (-80°C) until processing. Stomach samples were collected, categorized by their fullness, and found to be mostly empty, precluding the morphological identification of diet.

In the laboratory, samples (approximately 1-2 g) were rinsed in deionized water to remove exogenous substances and parasites (Nematoda spp.) removed if present on the liver. Samples were subsequently freeze-dried and manually crushed and homogenised using a pestle. Due to excess lipid content, liver samples underwent lipid extraction by sonication for 10 minutes in a 2:1 dichloromethane/methanol solution, repeated a minimum of three times until the solution remained clear. Residual solvents were then evaporated overnight prior to sample storage at ambient room temperature. This method was chosen given that it reduces lipid bias in δ^13^C, while not impacting δ^15^N (Andrews et al., 2023; Søreide et al., 2006), that can otherwise be a consequence of lipid extraction (Svensson et al., 2016).

### 2.2. Stable isotope analyses

To determine carbon, nitrogen and sulphur isotopic values, freeze-dried muscle and liver (0.9 to 1.2 mg respectively) were analysed in 20% duplicate using a Delta V Advantage continuous-flow IRMS coupled via a ConfloIV to an IsoLink elemental analyser (Thermo Scientific, Germany) at SUERC (East Kilbride, U.K.) as described in (Sayle et al., 2019). The obtained values were corrected from the isotopic ratio of the international standards, Vienna Pee Dee Belemnite (VPDB) for carbon, air (AIR) for nitrogen, and Vienna Cañon-Diablo Troilite (VCDT) for sulphur, using the δ (‰) notation.

Uncertainties on the measurements were calculated by combining the standard deviation (SDs) of the sample replicates and those of International Atomic Energy Agency (IAEA) reference material according to (Sayle et al., 2019). The International Atomic Energy Agency (IAEA) reference materials USGS40 (L-glutamic acid, δ^13^C_VPDB_ = –26.39 ± 0.04‰, δ^15^N_AIR_ = –4.52 ± 0.06‰) and USGS41a (L-glutamic acid, δ^13^C_VPDB_ = 36.55 ± 0.08‰, δ^15^N_AIR_ = 47.55 ± 0.15‰) were used to normalise δ^13^C and δ^15^N values. An in-house standard (5-SAAG, δ^34^S_VCTD_ = –12.86 ± 0.20‰) that is calibrated to the International Atomic Energy Agency (IAEA) reference materials IAEA-S-2 (silver sulfide, δ^34^S_VCTD_ = 22.62 ± 0.08‰) and IAEA-S-3 (silver sulfide, δ^34^S_VCTD_ = –32.49 ± 0.08‰) and the Iso-Analytical standard IA-R069 (tuna protein: δ^34^S_VCTD_ = 19.21 ± 0.22‰) were used to normalise δ^34^S values. Normalisation was checked using either the Elemental Microanalysis standards B2219 (Coldwater fish gelatin: δ^13^C_VPDB_ = –16.33 ± 0.10‰, δ^15^N_AIR_ = 14.71 ± 0.14‰, and δ^34^S_VCTD_ = 17.05 ± 0.07‰), B2215 (Fish gelatin: δ^13^C_VPDB_ = –22.92 ± 0.10‰, δ^15^N_AIR_ = 4.26 ± 0.12‰, and δ^34^S_VCTD_ = 1.21 ± 0.24‰), B2222 (Bovine gelatin: δ^13^C_VPDB_ = –11.11 ± 0.09‰, δ^15^N_AIR_ = 7.54 ± 0.12‰, and δ^34^S_VCTD_ = 6.79 ± 0.08‰), or the IAEA reference material USGS88 (marine collagen: δ^13^C_VPDB_ = –16.06 ± 0.07‰, δ^15^N_AIR_ = 14.96 ± 0.14‰, and δ^34^S_VCTD_ = 17.10 ± 0.44‰),

Precision was determined to be ± 0.15‰ for δ^13^C, ± 0.2‰ for δ^15^N and ± 0.4‰ for δ^34^S and is based on long-term repeated measurements of the well-characterised Elemental Microanalysis IRMS fish gelatin standard B2215. The mean uncertainty between replicates across all muscle measurements (35 of 178) was 0.13 ± 0.18‰ for δ^13^C, 0.05 ± 0.09‰ for δ^15^N, and 0.19 ± 0.21‰ for δ^34^S. The mean uncertainty across all liver measurements (36 of 179) was 0.14 ± 0.14‰ for δ^13^C, 0.11 ± 0.31‰ for δ^15^N, and 0.23 ± 0.37‰ for δ^34^S.

The quality of δ^13^C values for the 178 muscle and 179 liver samples were assessed and controlled using C:N ratios given that an excess of lipids (higher C:N ratio = greater lipid proportion) can downwardly bias δ^13^C values, typically for samples where C:N ratios >3.5 (Skinner et al., 2016). Across all samples and sites for each species, muscle C:N ratios were between 3.4-3.9 (mean 3.6) (Table S1) and there was no significant decrease of δ^13^C muscle values with C:N ratios (Pearson’s >0.05). Across all samples and sites for each species, liver C:N ratios were between 3.6-8.7 (mean 4.7), resulting in a significant decrease of δ^13^C liver values with C:N ratios (Pearson’s <0.05). We mathematically corrected both muscle and liver δ^13^C values following (Hoffman and Sutton, 2010).

δ^13^Ccorrected = δ^13^Cbulk + (−6.39‰ * (3.76 − C:Nbulk))/C:Nbulk where δ^13^Cbulk and C:Nbulk represent the uncorrected δ^13^C value and [C] to [N] ratio from our stable isotope analyses and −6.39 ‰ and 3.76 are mean estimates of Δδ^13^Clipid and C:Nprotein derived from a dataset of 48 fish species (Hoffman and Sutton, 2010). δ^15^N and δ^34^S values were quality controlled by confirming that there were no significant relationships (Pearson’s >0.05) with C:Nbulk across all samples. δ^13^Ccorrected will hereafter be referred to as δ^13^C.

### 2.3 Statistical analyses

We tested statistical pairwise differences between species and sites using non-exact pairwise Wilcoxon tests in R, adjusting *p* values for false discovery rate as per (Benjamini and Hochberg, 1995). To estimate the probability of each site being found within the same niche, for each species, we applied the overlap function using default settings and 10,000 iterations in the R package nicheROVER (Swanson et al., 2015). We tested the relationship between Total Length (TL) and each isotope using the lm linear regression function in R, for each site and species. Three ling samples that were size outliers (<55 cm and >150 cm) were excluded from these analyses. A base10 log-linear model was applied to TL values as per (Nakazawa et al., 2010) and (Jennings, 2005). Relationships between TL and Fulton’s condition were similarly tested. Finally, we tested the statistical differences between muscle and liver isotope values using exact pairwise Wilcoxon tests, adjusting *p* values as previous.

## 3. Results

A total of 178 muscle and 179 liver samples were analysed for δ^13^C, δ^15^N, and δ^34^S stable isotopes. δ^13^C values ranged from -21.6 to -16.6‰, δ^15^N values ranged from 10.9 to 15.6‰, and δ^34^S values ranged from 17.1 to 21.4‰. Variations in species isotope values were found i.e. lower δ^15^N values for saithe compared to tusk and ling, indicating saithe fed lower down the food web (Figure 2A). Studying muscle, lower δ^13^C values were found for saithe suggesting that saithe fed on a greater proportion of pelagic prey and tusk had higher δ^13^C values than ling suggesting that tusk fed the most benthic and ling was an intermediate (Figure 2A,3A). Finally, δ^34^S values were higher for saithe and ling compared to tusk, suggesting that tusk fed more offshore than saithe and ling.

**Figure 2.**
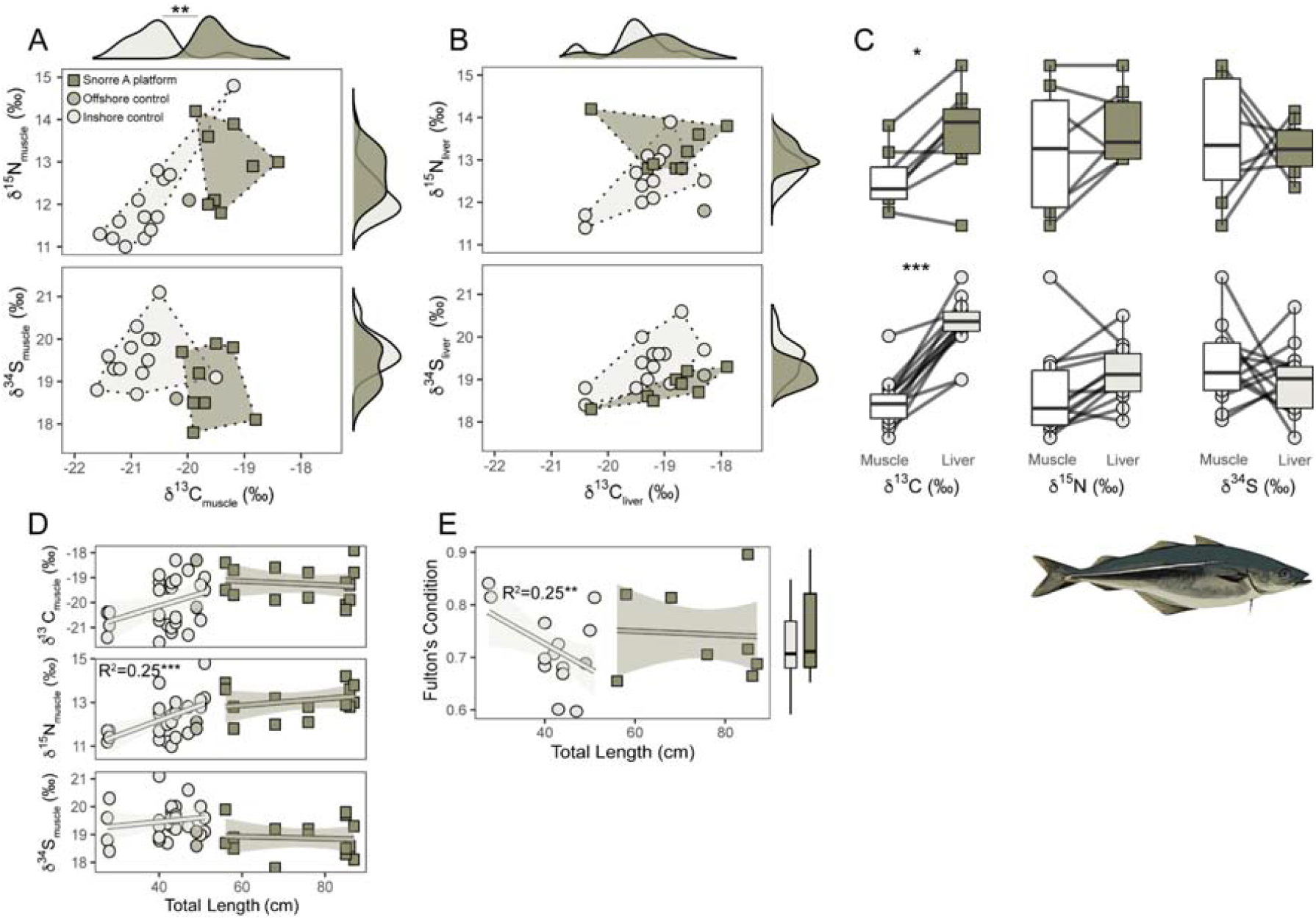
δ^13^C, δ^15^N and δ^34^S isotope measurements on saithe (*Pollacius virens*) illustrated using scatterplots of muscle (A) and liver (B) tissue at each of the sampling sites: Offshore and Inshore control sites (circles) and Snorre A (square). Convex hull total areas are shown for each site as dashed lines and density distributions are shown for each isotope with significance between groups tested by non-rank paired Wilcoxon tests. Boxplots (C) show differences between muscle and liver δ^13^C, δ^15^N and δ^34^S isotope measurements with grey lines joining pairs of samples from the same specimen and significance between groups as tested by ranked paired Wilcoxon tests. Relationships between total length (TL) and muscle isotope ratios (D) are shown using scatterplots and a lm smooth estimated using ggplot geom_smooth where shading indicates 95% CIs. Relationships were tested using linear regressions after TL was log10 transformed, where the regression coefficient (*R*2) and significance were calculated. Regressions (D) and testing between sites (C, E) excluded the offshore control site (green circles) due to low sample size (n=2). Relationships between Fulton’s Condition Index and TL are similarly shown where boxplots show differences between sample sites (E). Boxplots show group medians, 25th and 75th percentile as outer edges and outliers illustrated outside of 95th percentiles (black whiskers). Significance is represented as ‘*’ <0.05, ‘**’ <0.01 and ‘***’ <0.001 and is not shown if ‘ns’ >0.05.

**Figure 3.**
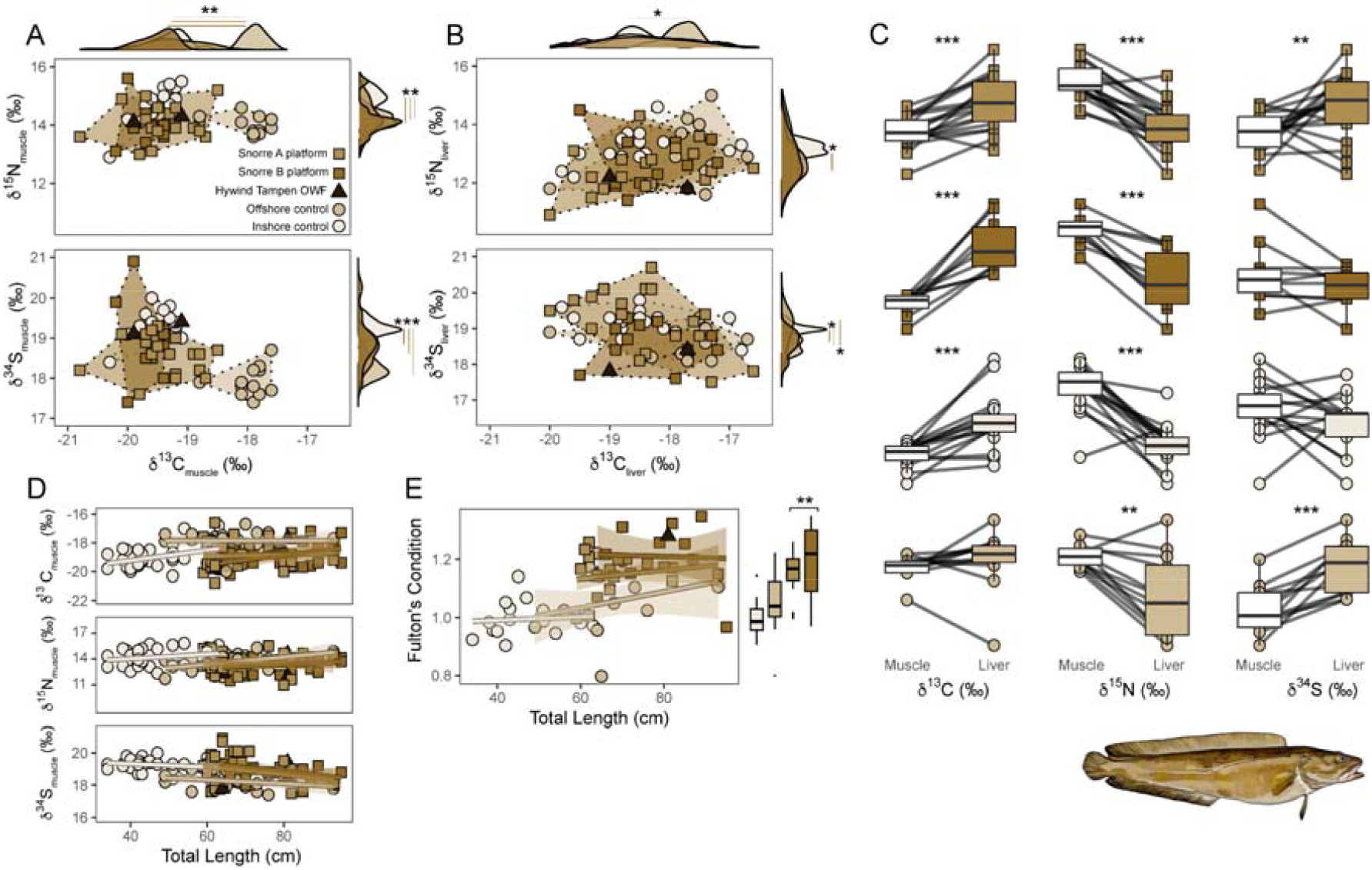
δ^13^C, δ^15^N and δ^34^S isotope measurements on tusk (Brosme brosme) illustrated using scatterplots of muscle (A) and liver (B) tissue at each of the sampling sites: Offshore and Inshore Control sites (circles), Hywind Tampen (triangle) and Snorre A and B (squares). Convex hull total areas are shown for each site as dashed lines and density distributions are shown for each isotope with significance between groups tested by non-rank paired Wilcoxon tests. Boxplots (C) show differences between muscle and liver δ^13^C, δ^15^N and δ^34^S isotope measurements with grey lines joining pairs of samples from the same specimen and significance between groups as tested by ranked paired Wilcoxon tests. Relationships between total length (TL) and isotope ratios (D) are shown using scatterplots and a lm smooth estimated using ggplot geom_smooth where shading indicates 95% CIs. Relationships were tested using linear regressions after TL was log10 transformed, where the regression coefficient (*R*2) and significance were calculated. Regressions (D) and testing between sites (C, E) excluded the Hywind Tampen (black triangles) site due to low sample size (n=2). Relationships between Fulton’s Condition Index and TL are similarly shown where boxplots show differences between sample sites (E). Boxplots show group medians, 25th and 75th percentile as outer edges and outliers illustrated outside of 95th percentiles (black whiskers). Significance is represented as ‘*’ <0.05, ‘**’ <0.01 and ‘***’ <0.001 and is not shown if ‘ns’ >0.05.

Overall, we found high niche overlap between offshore energy and control sites for each species and few consistently statistically significant differences between sites for each species when assessing muscle or liver.

### 3.1 Intraspecific patterns in Saithe

Saithe at Snorre A had higher but not significant δ^15^N values, significantly lower δ^13^C values (Wilcoxons p<0.01), and higher but not significant δ^34^S values (Figure 2A). Relatively low NicheRover overlap probabilities (18-21% for muscle, liver 13-38%, Figure S1A) were found between sites.

Significant differences between sites for saithe δ^13^C were not observed when studying liver measurements, and differences in δ^15^N decreased (Figure 2B). Comparing muscle and liver (Figure 2C), δ^13^C converged between sites, increasing to a greater extent between muscle and liver for the inshore control site than Snorre A (Wilcoxons p<0.001, p<0.05, respectively), while δ^15^N increased and δ^34^S decreased in liver, both non-significantly. Between samples, δ^13^C increased (non-significantly) with size, as did δ^15^N (inshore control: linear regression, p<0.001) while δ^34^S decreased (non-significantly, Figure 2D). Fulton’s condition did not exhibit a clear relationship with size despite inshore control saithe significantly decreasing with size (Figure 2E). Overall, saithe condition did not significantly differ between Snorre A and the inshore control (Figure 2E).

### 3.2 Intraspecific patterns in tusk

Tusk at the offshore control site had higher δ^13^C values than offshore energy and inshore control sites (Wilcoxons, p<0.01), while tusk at the inshore control site had significantly higher (Wilcoxons, p<0.01) δ^15^N values and significantly lower (Wilcoxons, p<0.001) δ^34^S values than other sites when assessing muscle measurements (Figure 3A). Sites shared relatively low overlap probabilities (0-27%, Figure S1B).

When assessing liver measurements, differences between sites in δ^13^C, δ^15^N and δ^34^S decreased (Wilcoxons, p<0.05, Figure 3B). Comparing muscle and liver (Figure 3C), δ^13^C converged between sites, increasing to a lesser extent between muscle and liver for the offshore control site (non-significant) than Snorre A, B and the inshore control (Wilcoxons, p<0.001), while δ^15^N significantly decreased in liver across all sites (Wilcoxons, p<0.01-0.001), and δ^34^S increased significantly in liver, at Snorre A and the offshore control site (Wilcoxons, p<0.01-0.001, Figure 3C). Between samples, no clear or significant relationships were found between isotope values and size (Figure 3D). Fulton’s condition increased with size in tusk, where larger tusk at Snorre A and Snorre B had significantly greater condition than the control sites (Figure 3E).

### 3.3 Intraspecific patterns in ling

Ling at the offshore control site had higher δ^13^C values than offshore energy sites (Wilcoxons, p<0.01), while ling at Highwind Tampen OWF had higher δ^13^C (Wilcoxons, p<0.01) values than Snorre A and B platforms, when assessing muscle measurements (Figure 4A). δ^15^N values at the offshore control site were significantly higher than Snorre A and B, as were those at Hywind Tampen (Wilcoxons, p<0.001). δ^34^S values at the offshore control site were significantly lower than Snorre A, B and Hywind Tampen (Wilcoxon, p<0.01) when assessing muscle measurements (Figure 4A). Sites shared high overlap probabilities (44-85%, Figure S1C).

**Figure 4.**
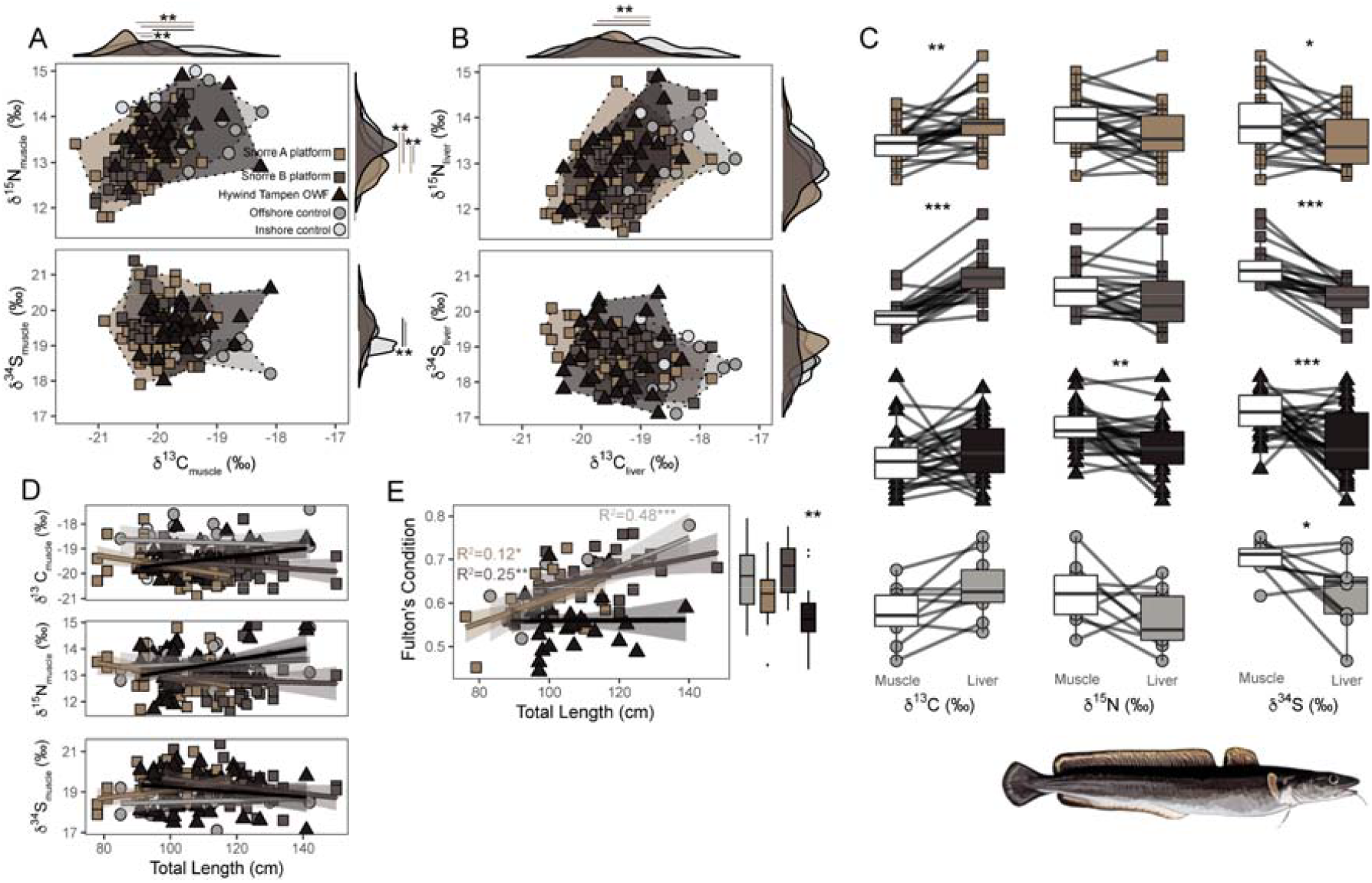
δ^13^C, δ^15^N and δ^34^S isotope measurements on ling (Molva molva) illustrated using scatterplots of muscle (A) and liver (B) tissue at each of the sampling sites: Offshore and Inshore Control sites (circles), Hywind Tampen (triangle) and Snorre A and B (squares). Convex hull total areas are shown for each site as dashed lines and density distributions are shown for each isotope with significance between groups tested by non-rank paired Wilcoxon tests. Boxplots (C) show differences between muscle and liver δ^13^C, δ^15^N and δ^34^S isotope measurements with grey lines joining pairs of samples from the same specimen and significance between groups as tested by ranked paired Wilcoxon tests. Relationships between total length (TL) and isotope ratios (D) are shown using scatterplots and a lm smooth estimated using ggplot geom_smooth where shading indicates 95% CIs. Relationships were tested using linear regressions after TL was log10 transformed, where the regression coefficient (*R*2) and significance were calculated. Regressions (D) and testing between sites (C, E) excluded the inshore control site (light grey circles) due to low sample size (n=2). Relationships between Fulton’s Condition Index and TL are similarly shown where boxplots show differences between sample sites (E). Boxplots show group medians, 25th and 75th percentile as outer edges and outliers illustrated outside of 95th percentiles (black whiskers). Significance is represented as ‘*’ <0.05, ‘**’ <0.01 and ‘***’ <0.001 and is not shown if ‘ns’ >0.05.

When assessing liver measurements, differences between sites in δ^15^N and δ^34^S decreased and only significant differences in δ^13^C values between the offshore control site and offshore energy sites remained (Wilcoxons, p<0.01, Figure 4B). Comparing muscle and liver (Figure 4C), δ^13^C converged between sites, increasing to a lesser extent between muscle and liver for the offshore control site (non-significant) than Snorre A, B (Wilcoxons, p<0.001), while δ^15^N decreased in liver across all sites, significantly at Hywind Tampen (Wilcoxons, p<0.01), and δ^34^S decreased significantly in liver, across all sites but to a lower degree at the offshore control site (Wilcoxons, p<0.05-0.001, Figure 4C). Between samples, no clear or significant relationships were found between isotope values and size (Figure 4D). Fulton’s condition increased with size in ling caught at Snorre A, B and the offshore control site (linear regressions, p<0.05-0.001), but not at Hywind Tampen which exhibited an overall significantly lower condition than the other three sites (Wilcoxons, p<0.01, Figure 4E).

## 4. Discussion

Our results identify few consistent differences between isotope values of codfishes caught at offshore energy structures and control sites indicating limited artificial reef effects of floating offshore structures on codfishes in the Norwegian North Sea. One reasonably predicts that artificial assemblages of fouling and aggregating fauna at offshore energy structures would comprise unique isotope values pertaining to trophic functional differences between native pelagic and soft-bottom fauna e.g. (Guerin 2009; van der Stap et al., 2016; Buyse et al., 2023) and that codfish diets significantly altered by such assemblages would engage in more resident behaviours than natural; as shown by a host of studies from the southern North Sea (Fujii and Jamieson, 2016; Lawrence et al., 2024; Reubens et al., 2013a), and that this in turn would alter stable isotope values of fish tissues. Instead, we find that δ13C isotope differences in saithe between the O&G and control sites were confounded by differences in body size, as was the greater condition of tusk at O&G sites. The combination of significantly lower condition and δ15N values of ling at the OWF site may conversely be explained by its reduced food availability given Hywind Tampen OWF was a new site established in 2022, but comparisons between longer-temporal perspective muscle and shorter-temporal perspective liver samples indicate δ15N values of ling reduced at all sites in the weeks prior to capture. Overall, our results support the idea that artificial reef effects are temporal phenomena not necessarily altering the diet of fishes, and that differences in reef effects between shallow and deep habitats and between fixed and floating technologies can be expected.

### Species- and site-specific-artificial reef effects

It remains challenging to conclude on the impact of artificial reef effects on fishes due to differences in effects depending on the functional ecology of each species, habitat and environment. For example, fishes that are ‘complex-bottom’ or reef fishes are likely to differ in responses than pelagic c.f. (Methratta and Dardick, 2019) depending upon niches provided by offshore energy structures that will be unique to the habitats they are installed in. Functional differences are likely to explain why studies on abundance e.g. (Bicknell et al., 2025) have found differences in effects between species and sites and this logically extends to diet effects (Buyse et al., 2023; Reubens et al., 2013b). In this study, we evaluated reef effects on three bentho-pelagic codfishes with varying functional ecology being a) saithe - the most pelagic and low trophic level (lowest δ^13^C and δ^15^N values) of the three, b) tusk - the most benthic (highest δ^13^C values) and c) ling - an intermediate that fed less offshore (lower δ^34^S values). Our stable isotope results match expectations of the three species’ life-histories; with saithe being the most long-ranging and surface-dwelling, tusk and ling being more sedentary, and ling diet comprising of greater proportions of fish than tusk (Bergstad, 1991; Tenningen et al. 2024).

Our stable isotope results indicate that saithe at Snorre A consumed a greater proportion of benthic, or deeper dwelling, and offshore prey than control sites despite the saithe diet comprising of a mix of bentho-pelagic prey at all sites. This aligns with expectations (and liver vs. muscle measurements) that older, larger saithe at Snorre A were more influenced by offshore, benthic prey at higher trophic levels. It is crucial to assess size effects on isotope values, condition, and differences between muscle and liver, which reflect seasonal and age patterns. Isotope differences between control and OWF sites may have been exaggerated in other studies, as differences are only visible when group means are taken and are confounded by size effects (Buyse et al., 2023). Additionally, our results suggest greater proportions of benthic prey for tusk at Snorre A and B, Hywind Tampen, and the Coastal control site, than the offshore control site, suggesting no significant differences in benthic versus pelagic foraging at offshore energy sites. Similarly, ling occupied a similar trophic niche at each of the sites. Our observation that differences in isotope values between sites decreased for each species when assessing liver compared to muscle, suggests that natural variation in diet over time was greater in muscle (representing a longer time average before capture) than liver (representing a shorter time average in the weeks prior to capture).

Given that some studies have found the opposite of our findings in the southern North Sea, reasoning may be one or a combination of a) physical characteristics of floating structures i.e. supporting less life than fixed ones, b) seasonal i.e. spring effects being limited compared to late summer when the majority of studies are conducted, or c) characteristics of the species studied/attracted and their habitats. Each of these factors are likely to impact the impact of artificial reef effects on fish populations and require disentangling through further study.

### Temporal relevance of species attraction

Generally studies on fish aggregation at offshore structures lack good temporal resolution; we compared values in short-lived liver and long-lived muscle tissue to overcome this challenge. Our findings support the idea that artificial reef effects of offshore energy structures may not be widespread across species and sites but also in time. The duration of impact has key consequences at the population and ecosystem level and our findings of limited artificial reef effects in the Norwegian North Sea are perhaps expected given that gill netting showed no increase in saithe, tusk or ling abundance around Hywind Tampen (Tenningen et al. 2024) but rod and line fishing closer to the structures (anchors) had the potential to target more resident individuals. Saithe and ling were common in the area prior to the establishment of the OWF and we cannot exclude the fact that differences, e.g. in ling condition at the OWF were not related to depth effects, despite depths between sites being similar, because of the temporal phenomenon of fish being diurnally or temporally attracted to energy structures (Fujii and Jamieson, 2016; Lawrence et al., 2024; Reubens et al., 2013a). In summary we advocate for improved temporal resolution in diet studies that cannot be achieved from snap-shots of stomach content analyses e.g. Tenningen et al. (2024) however multi-tissue isotope analyses as herein carry strong advantages as shown that avoid the overinterpretation of small differences between sites when using single-tissue measurements reported in other studies (Buyse et al., 2023).

### Limitations of the approach

We find it unlikely that a lack of consistent differences between sites was a result of the resolution offered by SIA because we found clear differences with ecological explanations between sample types of muscle and liver, and between species. SIA has been consistently shown to provide valuable insights into the dietary habits of fish (Amiraux et al., 2023; Mavraki et al., 2021; Sigman et al., 2009) but we acknowledge every approach has its limitations. One way to improve the results (at greater cost) could be to employ compound specific stable isotope analyses that allow for disentangling trophic vs. source (location) contributions of stable isotope signatures that we could not explore here. In addition, one could explore SIA of prey items to validate expectations of isotopic differences between natural and artificial fouling and attracted organisms. Other proxies e.g. fatty acid (FA) profiles could also be used that may be more distinct compared to stable isotope signatures (Plumlee et al., 2021; Rooker et al., 2006). These more complex approaches may be more effective in detecting dietary shifts, particularly when trophic discreteness in consumers is low, as seen in opportunistic or omnivorous feeding strategies (Persson et al., 1996).

Methodological enhancements could also be useful to account for size effects but here we find no significant relationships between stable isotope values and total lengths, except for saithe that we conclude drive its isotopic differentiation between O&G and control sites. Additionally, the potential impact of lipid extractions on δ15N and δ34S values cannot be entirely excluded. Although lipid extractions theoretically bias nitrogen values upwards, our findings indicate lower nitrogen levels in fatty liver tissues with higher C:N ratios (Ouellet et al., 2024). Furthermore, lipid extractions were corrected for carbon, and no significant differences in suphur values were observed between groups. While lipid extractions are expected to decrease sulphur values, this effect is not consistent across sample types and remains underexplored (Riverón et al., 2022). Therefore, while stable isotope analyses are robust, these factors should be considered when interpreting the results.

### Consequences for management and conservation

Clearly, year-round studies using a greater number of sites are required to draw more comprehensive conclusions. Nonetheless our results highlight that fish presence at offshore floating sites does not always result in enhanced secondary production. Further study is needed to determine the impacts of fish attraction (Methratta and Dardick, 2019), whether food provision occurs or not. Artificial reefs and additional efforts to ‘increase biodiversity’ through improper decommissioning of O&G and the ‘nature-inclusive design’ of OWFs e.g. micro-siting and add-on installations like ‘fish hotels’ may result in unwanted and negative impacts, such as ecological-traps that hinder ecosystem restoration (Methratta and Dardick, 2019; Pardo et al., 2023; Reubens et al., 2013b; Rouse et al., 2019). Challenges for management arise given that impacts of fish attraction may be compounded by intensive fishing that has occurred over centuries in the North Sea, and that population and ecosystem vulnerability is not represented by vulnerability frameworks e.g. IUCN red list (Bennun et al., 2024). Similar to fishery-exclusion effects inside OWFs and safety zones of O&G structures, the aggregation of fish to offshore structures is not entirely negative or positive (Coates et al., 2016). But as shown from years of Marine Protected Area (MPA) research, fishing at aggregation sites and site edges on greater densities of aggregating fish (Methratta and Dardick, 2019) has the potential to be more efficient than otherwise (Grossman et al., 1997; Lawrence et al., 2024), requiring consideration by policy-makers and offshore energy developers.

In conclusion, our study suggests that bentho-pelagic codfish diet and habitat use is not significantly or consistently altered by floating offshore energy structures in the Norwegian North Sea over periods of weeks, contrasting to limited findings from shallow southern North Sea habitats (Buyse et al., 2023; Mavraki et al., 2021). However, we found evidence for reduced condition and δ15N values of ling at the OWF site that may indicate a lack of food provision and potentially, fitness. With this study we show how stable isotope and simple condition analyses can be useful indicators of offshore wind farm effects that can enhance offshore monitoring programs.

## CRediT authorship contribution statement

AJA and SB designed the study. SB collected samples for analysis. AJA and SB conducted the laboratory work. AJA analysed the data. AJA wrote the manuscript. Both authors revised the manuscript.

## Acknowledgements

We thank Equinor ASA representing Offshore Norway for part-funding the work and Rolf Sundt and Lars-Petter Myhre for organising the sampling campaigns and catching the fish. We thank the Water Column Monitoring Consortia and associated researchers at NIVA, the Institute of Marine Research and NORCE for sample collection and Kerry Sayle for laboratory assistance. This work was funded by the Research Council of Norway (contract number 342628/L10) and Equinor ASA through the Water Column Monitoring 2024 Campaign as part of regulatory requirements to Miljødirekoratet (Norwegian Environment Agency).

## Data availability

Raw isotope data is found in the attached Appendix A: Supplementary data Table S1 along with meta-data for each sample.

