## Appendix B for "Stable isotope insights into artificial reef effects of floating offshore energy structures in Norwegian North Sea codfishes"

*Supplementary analysis figure*

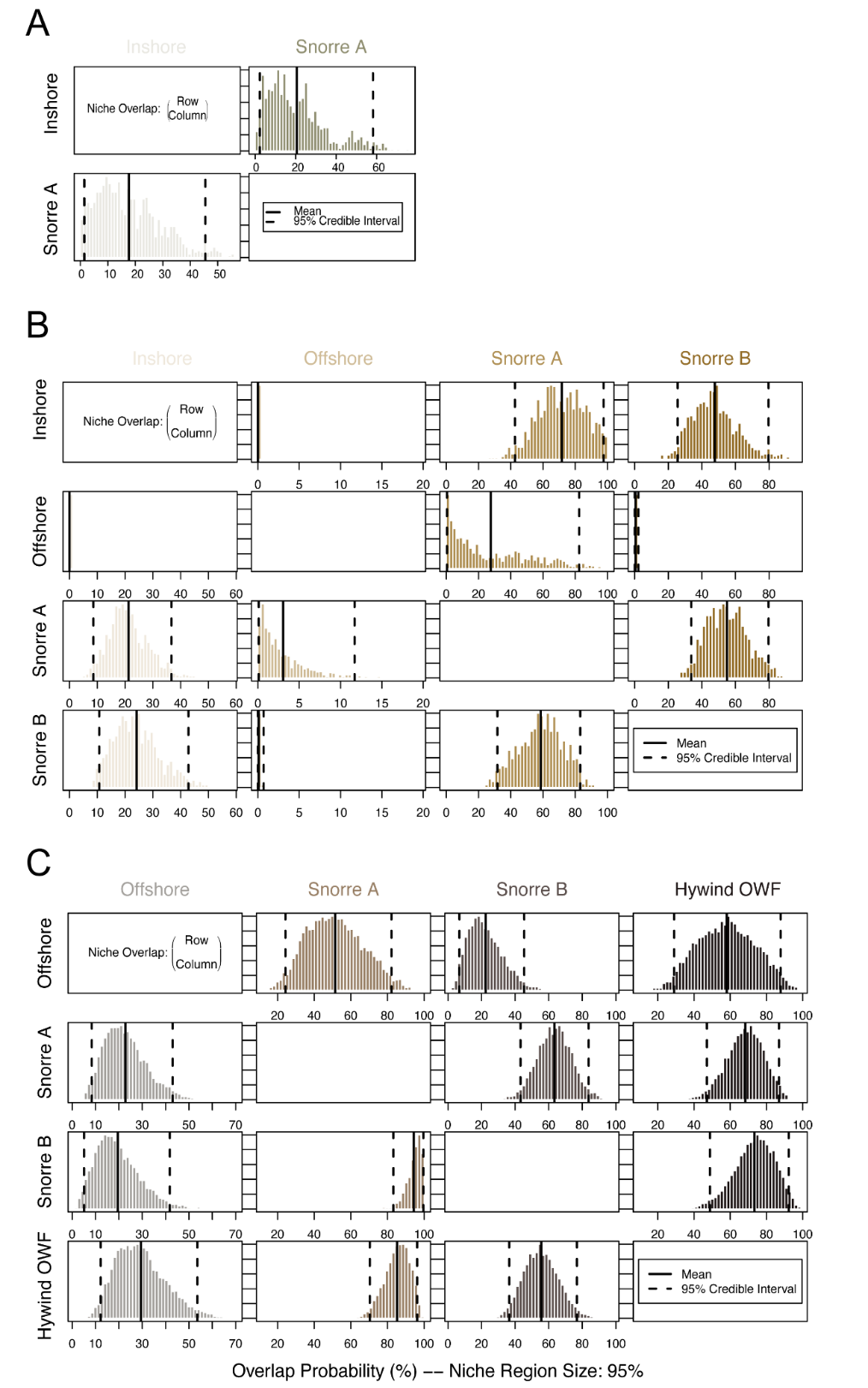
**Figure S1.** Overlap probabilities of each a priori defined site for saithe (A), tusk (B) and ling (C), showing the mean and 95% CI for each group pairing as estimated using nicheROVER using 10,000 permutations.
